# Reconstruction of 3-dimensional tissue organization at the single-cell resolution

**DOI:** 10.1101/2023.01.04.522502

**Authors:** Yuheng Fu, Arpan Das, Dongmei Wang, Rosemary Braun, Rui Yi

## Abstract

Recent advances in spatial transcriptomics (ST) have allowed for the mapping of tissue heterogeneity, but this technique lacks the resolution to investigate gene expression patterns, cell-cell communications and tissue organization at the single-cell resolution. ST data contains a mixed transcriptome from multiple heterogeneous cells, and current methods predict two-dimensional (2D) coordinates for individual cells within a predetermined space, making it difficult to reconstruct and study three-dimensional (3D) tissue organization. Here we present a new computational method called scHolography that uses deep learning to map single-cell transcriptome data to 3D space. Unlike existing methods, which generate a projection between transcriptome data and 2D spatial coordinates, scHolography uses neural networks to create a high-dimensional transcriptome-to-space map that preserves the distance information between cells, allowing for the construction of a cell-cell proximity matrix beyond the 2D ST scaffold. Furthermore, the neighboring cell profile of a given cell type can be extracted to study spatial cell heterogeneity. We apply scHolography to human skin, human skin cancer and mouse brain datasets, providing new insights into gene expression patterns, cell-cell interactions and spatial microenvironment. Together, scHolography offers a computational solution for digitizing transcriptome and spatial information into high-dimensional data for neural network-based mapping and the reconstruction of 3D tissue organization at the single-cell resolution.

## Introduction

The cell is the basic building block of life. Tissues are composed of many heterogeneous cells, usually numbering in the millions or billions. Each cell has its own location and performs specific functions that contribute to the physiological function of the tissue. These functions can include adhesion, sensing the environment, and communication with other cells. The expression of genes within a cell determines not only its identity but also its ability to interact with neighboring cells. This relationship between gene expression and cell localization and tissue architecture has been supported by genetic studies that have shown that manipulating gene expression can cause reproducible structural changes in tissues during development and homeostasis. However, it is difficult to map individual cells to 3D space and reconstruct the organization of tissues, based on their gene expression patterns^1–3^. The development of single-cell RNA sequencing (scRNAseq) has permitted more accurate measurement of the transcriptome at the single-cell level^4^. More recently, spatial transcriptomic (ST) platforms have been developed to measure the transcriptome of localized regions. However, the resolution of ST is limited by the size of the micropatterned pixels, which are usually 10-100 μm in diameter and capture a mixture of transcriptomes from multiple cells within a pixel. As a result, the single-cell resolution ST has yet to be established^1–3^. Computational methods, including cell-type deconvolution of spatial pixels such as RCTD^5^ and single-cell spatial charting methods such as CellTrek^6^, have been developed to enhance the resolution of ST. However, these methods acquire the spatial information of ST pixels as the 2D registration, which is dependent upon the sectioning angle of the reference slide. Furthermore, single cells are usually mapped back to 2D spatial positions constrained by the reference slide, which can limit the utility to identify cell neighbors and study spatially dynamic gene expression patterns.

In this study, we aim to map single cells and their associated transcriptome to specific locations in 3D space in order to reconstruct tissue organization and study the transcriptomic dynamics of the reconstructed tissue microenvironment. To address the limitations of current ST and computational methods, we have developed a new computational framework called scHolography. Our approach is based on three concepts. First, we reason that a distributed description of a spatial location, based on the distance between each pixel and all other pixels, can more accurately define the location of a pixel than 2D coordinates alone. This inter-pixel spatial information can better capture the intrinsic organizing principles of the tissue, regardless of the sectioning angle or slide orientation. Second, to establish an accurate transcriptome-to-space (T2S) projection, we treat the spatial-information components (SICs) as a high-dimensional dataset that describes the spatial organization of the tissue. Finally, we use neural networks to learn the T2S transformation and implement the Gale-Shapley algorithm to identify stable-matching neighbors (SMNs), which assigns single cells and their associated transcriptome to unique spatial locations. This approach allows us to improve spatial resolution from a large spatial pixel to the single-cell level without the need for cell-type deconvolution. Based on these principles, we have developed scHolography and applied it to human foreskin samples, as well as a recently published dataset of human skin cancer ST samples^7^ and a well-studied mouse brain ST dataset from 10X Genomics Visium. Our results demonstrate the accuracy of scHolography for *de novo* 3D tissue reconstruction, and highlight its ability to identify the profiles of neighboring cells of any given type, investigate spatial cell heterogeneity within tissues, and identify differential gene expression patterns across a defined space.

## Results

### scHolography learns inter-pixel distance and reconstructs tissue organization

The scHolography workflow aims to resolve the spatial dynamics of tissue at the single-cell resolution. One of the major goals of scHolography is to establish the transcriptome-to-space (T2S) projection, which maps a defined transcriptome to a spatial location within the tissue. While it is widely appreciated that scRNAseq accurately measures the transcriptome and defines cellular state^9^, it remains unclear which parameters could be used to define the spatial identity of the cell. Because no two pieces of tissue sections are the same, and x- and y-coordinates from each section also depend on arbitrary features such as tissue orientation and sectioning angle, 2D coordinates of cells/pixels only capture limited information for the spatial identity of cells/pixels in the ST dataset. We reason that the spatial identity of individual cells or pixels, in the case of spot-based ST platform, should collectively reflect cell-cell interactions and 3D tissue organization globally. Therefore, the spatial identity of a given cell/pixel should be more accurately determined by the measurement of the distance between this cell/pixel to all other cells/pixels within the tissue, rather than relying solely on its 2D coordinates.

To develop scHolography, we acquire readily available 2D spatial registration from the 10x Visium platform and generate a high-dimensional spatial dataset by computing pairwise pixel-pixel distances from 2D ST registration. To generate the pixel-pixel distance matrix, each pixel is considered as a dimension, and distances from this pixel to all others are the measurement for the dimension of the pixel. Principal component analysis (PCA) is then performed on the distance matrix to select top-ranked PCs and their corresponding values for downstream inferences. We name these top-ranked PCs as spatial-information components (SICs) (Fig. 1a and Extended Data fig. 1a-j). To establish the transcriptome-to-space (T2S) projection, scHolography takes the spatial (ST) RNAseq and scRNAseq data, obtained from the same or similar samples, as input (Fig. 1a Input Data). To prepare data for model training, ST and SC expression data are first integrated into the shared manifold, and SIC values for each ST pixel are defined, as described above (Fig. 1a Data Preparation, see Methods).

**Figure 1.**
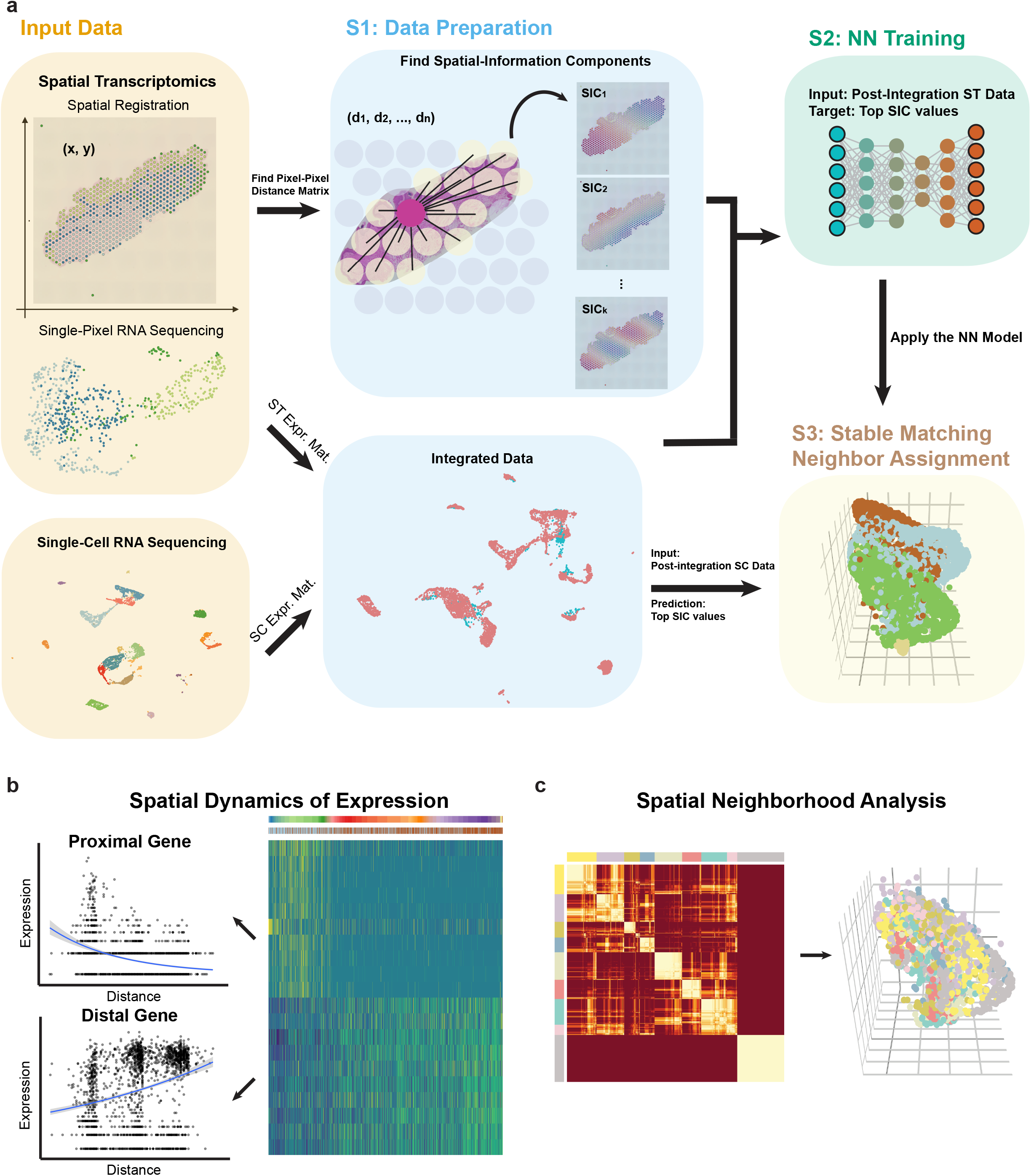
Overview of the scHolography workflow. **a,** Three steps of the scHolography workflow. (1) scHolography takes in ST and SC expression data and ST 2D spatial registration data. Spatial-information components (SICs) are defined for the spatial registration data. ST and SC expression data are integrated. (2) Neural networks are trained with post-integration ST data as input and top SIC values as the target. (3) The trained neural networks are applied to post-integration SC data to predict top SIC values for SC. SIC values are referenced to infer cell-cell affinity and construct the stable matching neighbor (SMN) graph. The graph is visualized in 3D. **b,** Based on inferred spatial distances among cells on the SMN graph, scHolography determines spatial dynamics of gene expression. The spatial gradient is defined as gene expression changes along the SMN distances from one cell population of interest to another. **c,** scHolography allows spatial neighborhood analysis. Cells are clustered according to their neighbor cell profile.

Next, scHolography trains neural networks to perform the T2S projection. Specifically, we use ST expression data as training input and SIC values as training targets for generating the T2S projection model (Fig. 1a NN training). The trained model is then applied to scRNAseq data to infer cell-cell affinity, a measurement for cell-cell virtual distance, from the predicted SIC values. The Gale-Shapley algorithm is then implemented to find stable-matching neighbors (SMNs) for each cell by using the cell-cell affinity matrix. Finally, scHolography reconstructs 3D tissue organization by projecting the cell-cell spatial connection with an undirected SMN graph, which could be visualized in 3D with the forced-directed Fruchterman-Reingold layout algorithm (Fig. 1a Stable Matching Neighbor Assignment).

In the reconstructed 3D tissue, each cell is assigned to a unique spatial location, and the distance between single cells is determined by the length of the shortest path connecting individual cells on the SMN graph. Thus, local and global tissue organization can be examined by ordering cells of interests based on their distances to any reference cell type and plotting the dynamics of gene expression patterns across inferred spatial organization of the tissue (Fig. 1b). Furthermore, cell heterogeneity can be studied by spatial organization, based on the first-degree neighbor profiles of any cell type (Fig. 1c), in addition to widely used transcriptomic clustering^10^. Collectively, scHolography reconstructs tissue organization in 3D, allows the identification of dynamic gene expression patterns across tissues, and determines spatial cell heterogeneity.

### scHolography recapitulates global and local spatial organization of human skin

We generated scRNAseq and ST datasets from human foreskin samples and examined the performance of scHolography. We generated ST datasets from 2 serial, sagittal sections from donor #1 (Fig. 2a and Extended Data fig. 2a). Our scRNAseq data, obtained from a different donor (donor #2), captured 6,425 cells with a mean depth of 136,235 reads/cell, and 5,450 cells passed our filtering with the Seurat package^10^. Unsupervised clustering identified major epithelial and dermal cell types. We also detected PECAM1+ endothelial cells, MGST1+ glandular epithelium, CD74+ immune cells, PROX1+ lymphatic endothelial cells, PMEL+ melanocytes, MPZ+ Schwann cells, and TAGLN+ smooth muscle cells (Fig. 2b). The human foreskin ST data (the serial section #1) from donor #1 captured 659 pixels with a median depth of 156,332 reads/pixel. By plotting with markers for major skin cell types, we confirmed that our ST data capture all major cell types in the skin, including epithelium, fibroblast, endothelial and smooth muscle cells (Fig. 2a and Extended Data fig. 2a-b).

**Figure 2.**
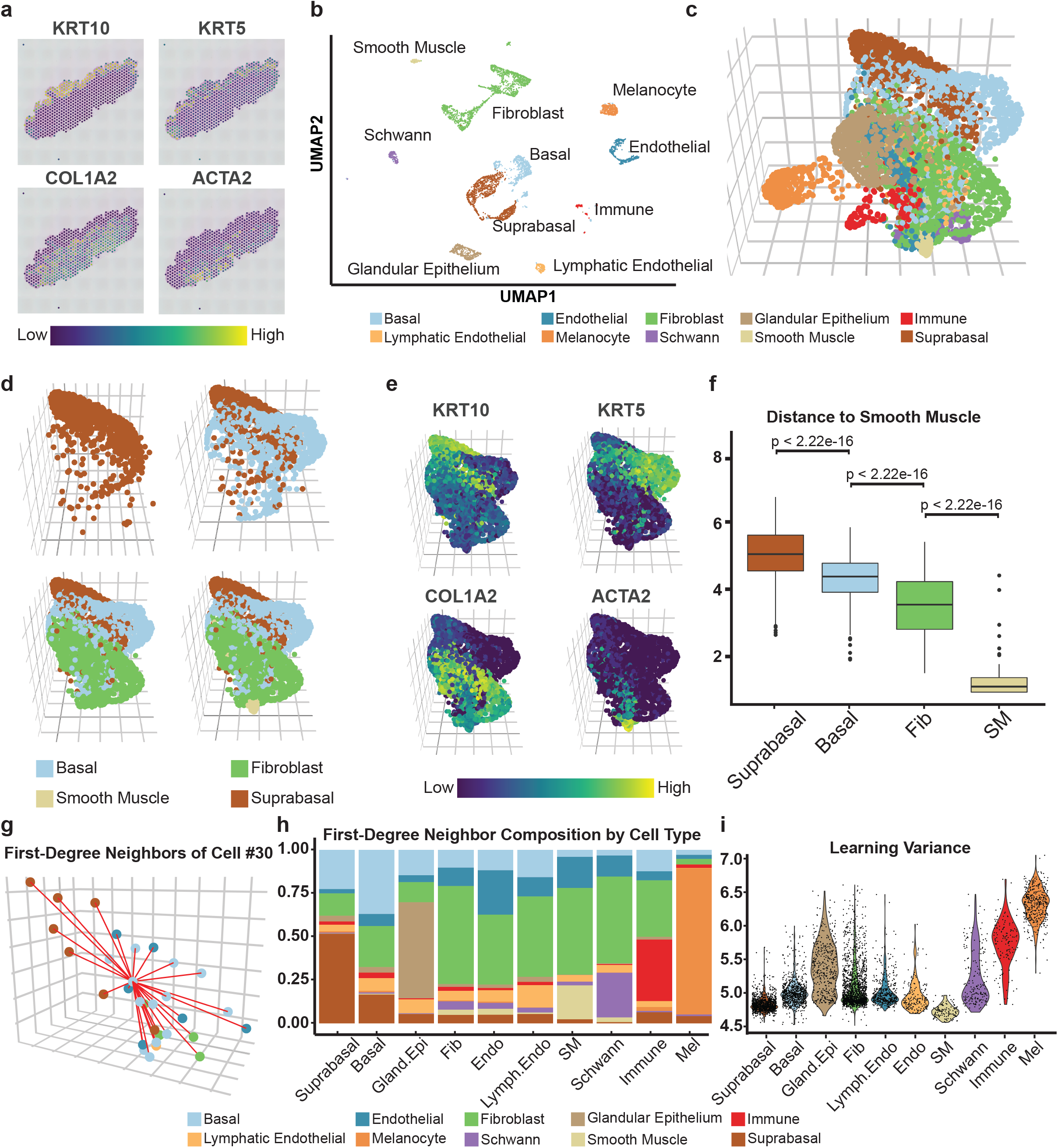
scHolography reconstructs the spatial organization of human foreskin. **a,** Spatial feature plots of markers for major cell types in Donor 1 slice 1 human foreskin ST data. *KRT10*, suprabasal cell marker; *KRT5*, basal cell marker; *COL1A2*, fibroblast marker; *ACTA2*, smooth muscle cell marker. **b,** UMAP plot of human foreskin scRNAseq data. **c,** 3D visualization of human foreskin spatial reconstruction by scHolography. **d,** scHolography 3D plot for 4 major cell types in the skin. **e,** scHolography 3D feature plot of marker genes for 4 major cell types. **f,** SMN distances between 4 major foreskin cell types to smooth muscle cells (Suprabasal cells n = 1120; Basal cells n = 808; Fibroblasts n = 1651; Smooth muscle cells n = 119). Boxplots show the median with interquartile ranges (IQRs) and whiskers extend to 1.5× IQR from the box. One-sided Wilcoxon tests are performed. **g,** scHolography 3D plot of Cell #30 and its first-degree neighbors. **h,** First-degree neighbor composition plot of major cell types in human foreskin. **i,** Violin plot of scHolography learning variance for each cell type in human foreskin.

To evaluate the performance of T2S projection and SMN assignment of our algorithm, we benchmarked scHolography against 2D spatial charting methods CellTrek and Seurat-SrtCT using the human scRNAseq and ST data. To compare the predicted tissue organization, we used the serial section #2 from donor #1 of human foreskin ST data as the ground truth (Extended Data fig. 2c-d) and reconstructed the serial section #2 by applying the model learned from the serial section #1 to the ST data of #2 (Extended Data fig. 2e-g). This allowed us to compare the reconstructed result to the experimentally determined result. We assessed the global prediction accuracy by calculating the pixel-by-pixel spearman correlation between reconstructed and experimentally determined coordinates (Extended Data fig. 2h, see Methods). We also evaluated the local accuracy by calculating the Jaccard similarity of the overlap in reconstructed and experimentally determined neighbors of pixels (Extended Data fig. 2i, see Methods). With both assessments, scHolography achieved the highest prediction accuracy (Mann Whitney Wilcoxon test, p<2.22e-16), significantly outperforming both CellTrek and Seurat-SrtCT. In addition, each pixel in scHolography reconstruction was uniquely assigned. In contrast, pixels in CellTrek reconstruction and, more prominently, in Seurat reconstruction had more overlapping, likely due to the inability to distinguish transcriptomic similar pixels.

Next, we applied scHolography to the scRNAseq data to reconstruct human foreskin at the single-cell level by applying the model learned from the serial section #1 as the spatial reference (Fig. 2c and Extended Data fig. 3a). scHolography reconstruction recapitulated stereotypical positions of major cell types, reflected by both cell type annotation and gene marker expression in the reconstructed 3D structure. For example, suprabasal epithelial cells, marked by KRT10^hi^ expression, were located at the outermost layer of the 3D structure, and KRT5^hi^ basal epithelial cells were located beneath the suprabasal cells and sandwiched between the suprabasal epithelial cells and dermal fibroblasts (Fig. 2d-e). The ACTA2^hi^ smooth muscle cells were located at the bottom of the reconstructed 3D tissue, consistent with stereotypical cell organization of the skin. In contrast, neither CellTrek nor Seurat-SrtCT was able to reconstruct 3D skin organization (Extended Data fig. 3b).

The quantitative information of cell-cell distance, SMN distance, is embedded in the prediction of scHolography, allowing for the study of tissue architecture based on spatial distance. This enabled us to analyze the distance between individual cell layers. As an example, we calculated the distance from suprabasal, basal, fibroblast and smooth muscle cells to smooth muscle cells (Fig. 2f). Not only were the differences highly significant between each cell type (Mann Whitney Wilcoxon test, p<2.22e-16) but also the spatial order agreed with stereotypical tissue organization such that suprabasal cells were furthest away from smooth muscle cells, followed by basal cells and fibroblasts (Fig. 2f). Furthermore, the SMN graph designates 30 stable-matching cells to a cell as its first-degree neighbors (Fig. 2g). We next determined the first-degree neighbor composition for each cell type in the skin by averaging 30 neighboring cell type information for all cells from each cell type (Fig. 2h and Extended Data table 1). For basal, suprabasal and glandular epithelial cells, the most abundant neighbors to each cell type were themselves as one may expect. However, fibroblasts often emerged as the most abundant neighbors for cell types that were localized in the dermis, including endothelial cells, lymphatic endothelial cells and Schwann cells. Therefore, scHolography allows quantitative analysis of cell heterogeneity based on their neighbor cell composition. Interestingly, we also noticed that each cell has a different matching stability for their assigned 3D location, likely due to cell migration or differences in cell types detected by scRNAseq and ST. We then computed a motility score, called learning variance, for each cell such that the confidence of the T2S projection and SMN assignment can be quantified. Melanocytes, immune cells and, to a lesser extent, glandular epithelial cells and Schwann cells, showed higher motility scores (Fig. 2i and Extended Data fig. 3c). In contrast, suprabasal and basal epithelial cells and smooth muscle cells showed the lowest motility scores.

### Spatially defined single-cell gene expression dynamics and cell heterogeneity in human skin

Having established the accuracy of scHolography in recapitulating cell organizations in 3D, we next investigated spatial dynamics of gene expression across multiple cell types in the human foreskin by using the *findGeneSpatialDynamics* function (see Methods). We first ordered cells from basal and suprabasal epithelial cell clusters according to their computed distance to the fibroblast clusters. As expected, basal cells were proximal to fibroblasts, whereas suprabasal cells were distal to fibroblasts (Fig. 3a). We then identified spatially variable genes and correlated their expression levels to the distance away from fibroblasts. For example, proliferation-related genes, such as CENPF, TOP2A, ASPM, and MKI67, were proximal to the fibroblast-basal boundary and showed a declining trend away from the boundary (Fig. 3b), reflecting the exit of the cell cycle when keratinocytes moved upward and differentiated^11,12^. Differentiation genes, such as KRT1 and KRTDAP, were distal to the boundary and showed an ascending trend as keratinocytes reached the suprabasal layer.

**Figure 3.**
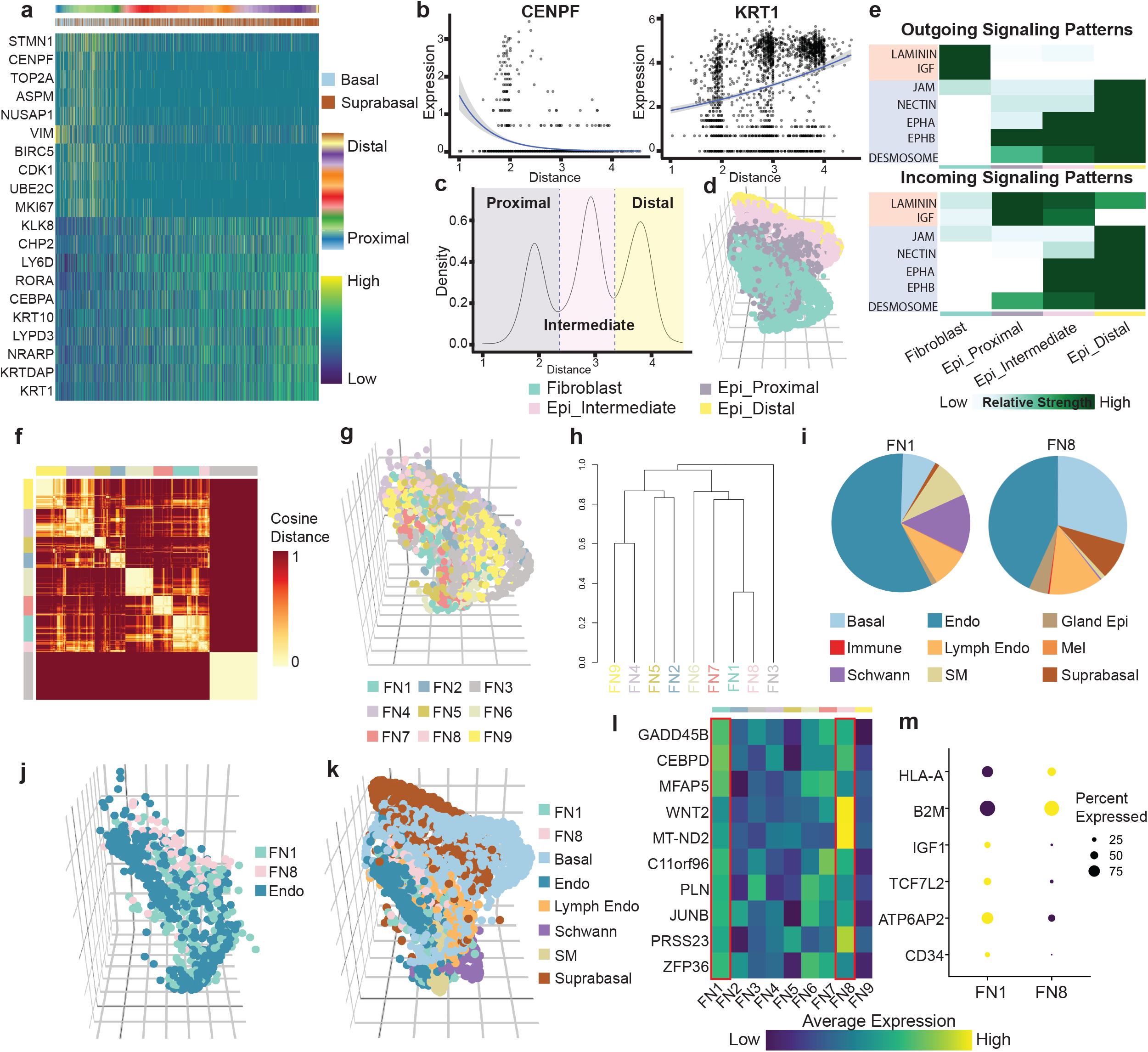
Single-cell gene expression dynamics and spatial cell heterogeneity in human skin. **a,** Expression heatmap of top ten spatially dynamic genes of human epithelial cells proximal (left) and distal (right) to fibroblasts. Poisson regression is performed to determine the significance. Epithelial cells are ordered, from left to right, in increasing SMN distance to fibroblasts. **b,** Expression-distance plot of *CENPF* (left, a proximal gene) and *KRT1* (right, a distal gene). 95% confidence intervals of Poisson regression are shown. **c,** Density plot of SMN distance of epithelial cells to fibroblasts. Epithelial cells are classified into proximal, intermediate, and distal epithelial cells by the distance percentile of 25% and 65%. **d,** scHolography 3D plot of epithelial cells and fibroblasts. **e,** Heatmap of relative outgoing (top) and incoming (bottom) strength of enriched signaling pathways predicted by CellChat. **f,** Spatial cell neighborhood analysis for fibroblasts. Nine distinct neighborhoods FN1-9 are identified based on the similarity of the first-degree neighbor cell composition. **g,** scHolography 3D plot of nine fibroblast spatial neighborhoods. **h,** Dendrogram based on the similarity of first-degree neighbor cell composition. **i,** Pie charts of non-fibroblast first-degree neighbor cell compositions for FN1 (left) and FN8 (right). **j,** scHolography 3D plot of FN1, FN8 and endothelial cells. **k,** scHolography 3D plot of FN1, FN8, basal, endothelial, lymphatic endothelial, Schwann, smooth muscle, and suprabasal cells. **l,** Heatmap of average expression levels of highly expressed genes in FN1 and FN8 (Mann Whitney Wilcoxon test, p<0.05). **m,** Feature dot plot of differentially expressed genes between FN1 and FN8 (Mann Whitney Wilcoxon test, p<0.05).

Interestingly, we observed a trimodal pattern of keratinocyte cell clustering patterns, based on their distance away from fibroblasts (Fig. 3c). We re-classified epithelial keratinocytes into 3 spatial clusters, epi_proximal, epi_intermediate and epi_distal cluster, according to the trimodal distance distribution, and visualized their 3D organization together with fibroblast cells (Fig. 3d). With these spatially defined clusters, we determined cell-cell communications between epithelial cells and fibroblasts by applying CellChat algorithm^13^. By using the inferred relative strength of signaling, we classified signaling into two types. Type 1 signaling, including laminin and IGF signaling, was originated from fibroblast and showed decreasing in-signal strength from proximal to distal epithelial cells, representing fibroblast-basal cell signaling events. Type 2 signaling, including NECTIN, EPHA and desmosome signaling, was originated from distal epithelial cells and showed a decreasing in-signal strength from distal to proximal epithelial cells, representing suprabasal-suprabasal cell signaling events. Notably, laminin and IGF signaling are related to basal cell adhesion to the basement membrane and basal cell proliferation^14^. In contrast, NECTIN, EPHA and desmosome, representing Type 2 signaling pathways, are related to tight junction and the initiation of epidermal differentiation^15,16^. These results from epidermal differentiation demonstrated the utility of scHolography for not only faithfully projecting single cells to reflect cell lineage differentiation but also identifying spatially dynamic gene expression patterns and signaling pathways.

We next performed quantitative analysis of spatial cell heterogeneity. We used dermal fibroblast cells as an example because of the high cell heterogeneity of these cells^17,18^. With the spatially resolved cell locations, we calculated cell type frequency-inverse cell frequency (CTF-ICF) to identify fibroblast subtypes with distinct first-degree neighbor cell compositions (Fig. 3f, see Methods). The composition of first-degree neighbor cells varied across spatial neighborhoods in the dermis, supporting the idea that different fibroblasts are located near different cell types, and nine distinct spatial neighborhoods for fibroblasts (FN1-9) were identified (Fig. 3g, Extended Data fig. 4a and Extended Data table 2). The hierarchical dendrogram of these nine clusters revealed similarities and differences among these nine spatial neighborhoods (Fig. 3h). Among them, FN1 and FN8 were the most similar pair of neighborhoods, and we further investigated their spatial features. The non-fibroblast first-degree neighbors of FN1 and FN8 were highly enriched for endothelial cells and lymphatic endothelial cells. However, FN1’s neighbors showed more abundant Schwann cells and smooth muscle cells, whereas FN8’s neighbors showed more basal and suprabasal epithelial cells (Fig. 3i). These observations suggested that FN1 fibroblasts were likely localized to deeper dermis and FN8 fibroblasts were localized to the surface of the skin. Indeed, visualization of FN1 and FN8 fibroblasts in 3D reconstruction corroborated these analyses (Fig. 3j-k).

To gain insights into differential gene expression of fibroblasts that was associated with different spatial neighbors, we determined enriched genes in FN1 and FN8 fibroblast populations. Notably, CEBPD, MFAP5 and WNT2, which have been shown to regulate fibroblast-endothelial cell interactions^19–21^, were enriched in both FN1 and FN8, consistent with their proximity to endothelial cells in spatial reconstruction (Fig. 3l). In addition to these commonly elevated genes, differential gene expression analysis identified CD34, ATP6AP2, TCF7L2 and IGF1 as highly enriched genes in FN1 fibroblasts and B2M and HLA-A, which are involved in MHC class I antigen presentation, as highly enriched genes in FN8 fibroblasts. Taken together, these results illustrate the utility of scHolography for not only reconstructing 3D tissue organization but also identifying spatial cell heterogeneity.

### Spatial reconstruction of human cutaneous squamous cell carcinoma

We next aimed to compare normal and diseased tissues and identify disease-associated spatial features. To achieve this goal, we applied scHolography to previously published human cutaneous squamous cell carcinoma (cSCC) datasets^7^, which contain both normal skin and cancerous regions (Fig. 4a). Furthermore, patient- and site-matched ST RNAseq and scRNAseq datasets were available for tissue reconstruction by scHolography (Fig. 4a-b). We applied scHolography, CellTrek and Seurat-SrtCT for tissue reconstruction. Only scHolography produced layered tissue patterns that were reminiscent of the reference tissue section with distinct normal and tumor regions (Fig. 4c and Extended Data fig. 5a-c).

**Figure 4.**
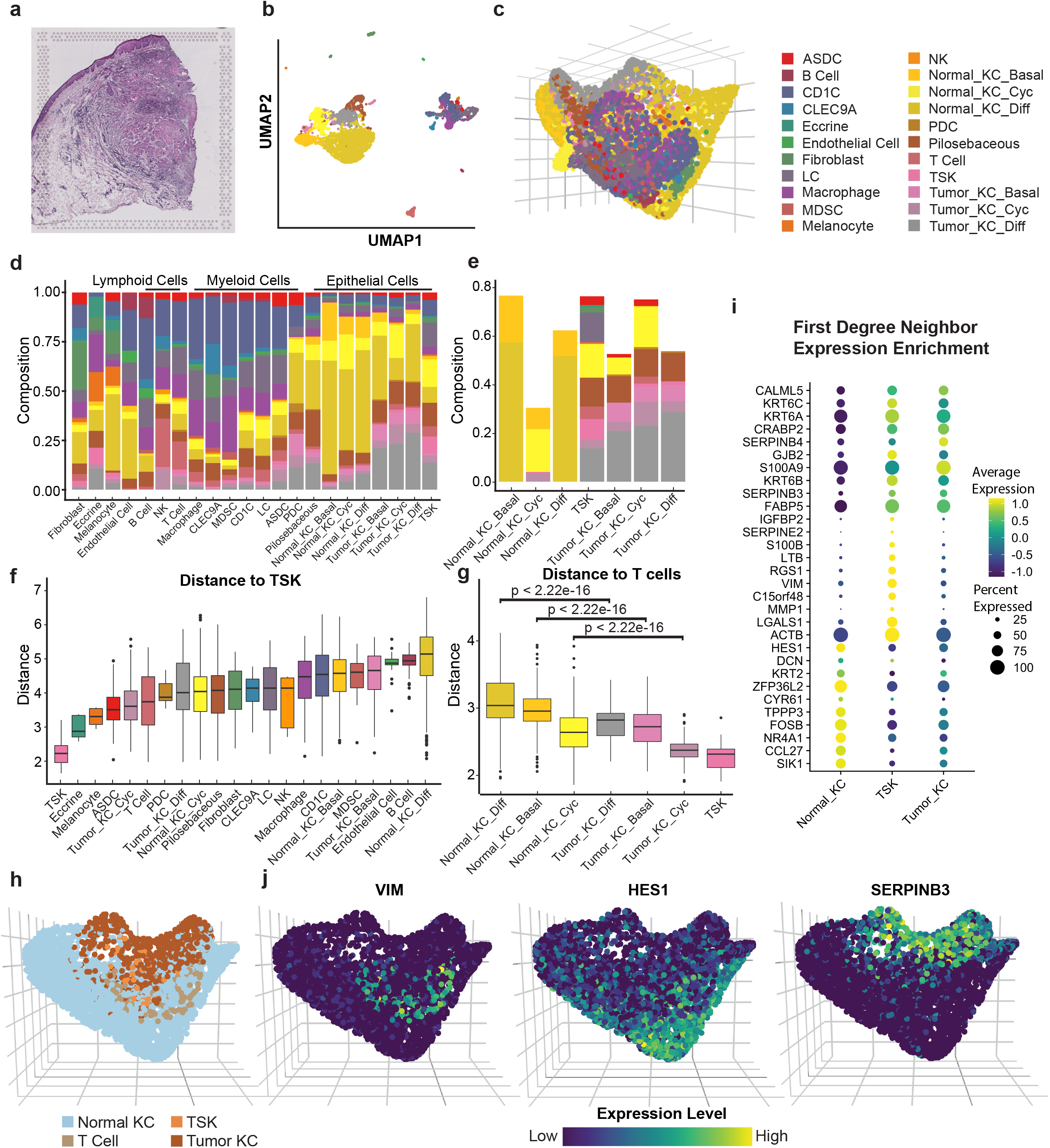
scHolography reconstructs the spatial organization of human cSCC. **a,** H&E image of Patient 6 rep 1 cSCC ST sample. **b,** UMAP plot of Patient 6 rep 1 scRNAseq data. **c,** 3D visualization of cSCC spatial reconstruction by scHolography. **d,** The first-degree neighbor composition plot for each cell type in cSCC. **e,** The first-degree neighbor composition plot for significantly enriched neighboring cell types of normal and tumor keratinocytes. **f,** SMN distances of major cell types to TSK cells by the order of increasing median distance (ASDC n = 70; B Cell n = 38; CD1C n = 595; CLEC9A n = 82; Eccrine cells n = 5; Endothelial Cells n = 23; Fibroblasts n = 114; LC n = 348; Mac cells n = 262; MDSC n = 18; Melanocytes n = 9; NK cells n = 5; Normal_KC_Basal n = 517; Normal_KC_Cyc n = 499; Normal_KC_Diff n = 2497; PDC n = 13; Pilosebaceous cells n = 385; T cells n = 128; TSK n = 34; Tumor_KC_Basal n = 116; Tumor_KC_Cyc n = 103; Tumor_KC_Diff n = 476). Boxplots show the median with interquartile ranges (IQRs) and whiskers extend to 1.5× IQR from the box. **g,** SMN distances of normal and tumor keratinocytes to T cells. One-sided Wilcoxon tests are performed to determine statistical significance. **h,** scHolography 3D plot of Normal KC, TSK, Tumor KC, and T cells. **i,** Feature dot plot of top ten significantly enriched genes among the first-degree neighbors of normal KC, including Normal_KC_Basal, Normal_KC_Cyc, Normal_KC_Diff, TSK, and tumor KC, including Tumor_KC_Basal, Tumor_KC_Cyc, Tumor_KC_Diff. Significances are determined by one-sided Wilcoxon tests. **j,** scHolography 3D feature plot of highly expressed genes among the first-degree neighbors of TSK (VIM), Normal KC (HES1), and Tumor KC (SERPINB3).

To benchmark the results, we used ST replicate #1 as the reference to learn the spatial information for the T2S projection. We next reconstructed ST replicate #2 in 3D and compared the accuracy of the T2S projection from scHolography, CellTrek and Seurat-SrtCT with the true spatial registration of replicate #2. For both global and local accuracy measurements, scHolography significantly outperformed the other two methods (Extended Data fig. 5d), mirroring the performance comparison from the human foreskin study.

To compare the spatial organization of normal skin and cSCC, we focused on the first-degree neighbor composition of each cell type. As expected, we observed a wide range of variation in the neighbor profile across different cell types (Fig. 4d, Extended Data fig. 5e and Extended Data table 3). Interestingly, cells from similar developmental origin, such as keratinocytes (KCs), myeloid cells and lymphoid cells, generally had more similar neighbor cell composition. In addition, immune cells, including both myeloid and lymphoid cells, generally had more complex neighbor cell compositions. Next, we turned to normal and tumor KCs to compare the neighbor profiles of normal vs diseased tissue regions. We also performed statistical test to identify the significantly enriched neighbor cell types for each KC subtype (Fig. 4e). Analysis of scRNAseq data identified basal, cycling and differentiated KCs in both normal and tumor regions. As described previously, tumor KCs also contained a unique cluster, named tumor-specific keratinocytes (TSKs)^7^. TSKs are enriched at the leading edge of tumor, and these cells demonstrate invasive and immunosuppressive features^7^. Interestingly, the first-degree neighbors of normal KCs, including basal, cycling and differentiated KCs, were largely composed of themselves or other normal KCs (Fig. 4e). Normal cycling KCs also had significant shares of TSK and tumor cycling KCs as their first-degree neighbors, revealing a higher degree of spatial heterogeneity of proliferative KCs compared with both basal and differentiating KCs. In sharp contrast, the first-degree neighbor profiles of tumor KCs were more complex with notable shares of immune cells, including T cells, plasmacytoid dendritic cells (pDCs) and AXL+SIGLEC6+ dendritic cells (ASDCs) (Fig. 4e). Pilosebaceous cells were also significantly enriched as the first-degree neighbors for all tumor KCs, suggesting a possibility that hair follicle stem cells serve as the cell of origin for tumor KCs^22^. Notably, TSKs showed the most diverse composition of the first-degree neighbors with the highest share of T cells, ASDCs, LCs, consistent with their location at the leading edge of the tumor.

Because of the key function of TSKs to tumorigenesis and invasion, we further investigated the microenvironment of TSKs. Leveraging the quantitative information of cell-cell distance embedded within scHolography, we determined the (averaged) distance between TSK cells and other cells. Besides themselves, TSKs were proximal to eccrine cells, melanocytes, AXL+SIGLEC6+ dendritic cells (ASDCs), cycling tumor KCs, T cells, and plasmacytoid dendritic cells (PDCs) (Fig. 4f). Interestingly, B cells and differentiated normal KCs were furthest away from TSKs. Because of the importance of T cell infiltration within tumor, we compared the distance from normal and tumor KC populations to T cells. While TSK showed the closest proximity to T cells among all KCs, all other tumor cell types, including cycling, basal and differentiated tumor KCs, were also significantly closer to T cells than their normal counterparts (Fig. 4g). Furthermore, cycling KCs, from both normal and tumor regions, were significantly closer to T cells than basal or differentiated KCs. This spatial proximity of TSKs and cycling cells to T cells suggests potential immune responses to the invasive and proliferating tumor cells, respectively, within the microenvironment of cSCC.

We next examined the 3D visualization of reconstructed tissue with a focus on normal KCs, tumor KCs, TSKs and T cells. Consistent with morphological findings^7^, TSKs were localized at the leading edge of the tumor and interacted closely with T cells (Fig. 4h). We next identified genes that were highly enriched within the first-degree neighbors of normal KCs, TSKs and tumor KCs. Consistent with distinct neighbor profiles for these cell types, we found differentially enriched genes within the neighbors of normal KCs, TSKs and tumor KCs (Fig. 4i). Specifically, ACTB, LGALS1, VIM and MMP1, which were associated with tumorigenesis and pro-progression^7^, were highly enriched in the neighbors of TSKs. Genes including FOSB, HES1 and ZFP36L2, which were associated with KC differentiation, were highly enriched in the neighbors of normal KCs, whereas SERPINB3 and SERPINB4, which are known for their roles in the initiation of the acute inflammatory response and as SCC antigen^23,24^, were enriched in the neighbors of tumor KCs (Fig. 4j). Taken together, scHolography reconstructs highly complex cSCC tissues and provides quantitative spatial information for investigating differential gene expression and studying tumor microenvironment.

### Spatial reconstruction of mouse brain

We applied scHolography to publicly available mouse brain data^8^ (Fig. 5a-b). Although the ST data were obtained from 2D brain slice (Fig. 5a), scHolography successfully reconstructed a well-defined tissue in 3D (Fig. 5c-d and Extended Data fig. 6a), characterized by the layered organization of GABAergic neurons, including Vip+, Pvalb+, Sst+, Sncg+ and Lamp5+ populations (Fig. 5e), glutamatergic neurons, and other non-neuronal cells, such as astrocytes and endothelial cells (Fig. 5f). Furthermore, the reconstruction of glutamatergic laminar excitatory neurons recapitulated the stereotypical organization, in the order of L2/3 intratelencephalic (IT), L4, L5 IT, L5 pyramidal tract (PT), L5/6 near-projecting (NP), L6 IT, L6 corticothalamic (CT) and L6b (Fig. 5g). We plotted the distance between the L2/3 IT neurons and all glutamatergic neurons on the scHolography SMN graph, and this quantification confirmed the layered organization of this region (Fig. 5h).

**Figure 5.**
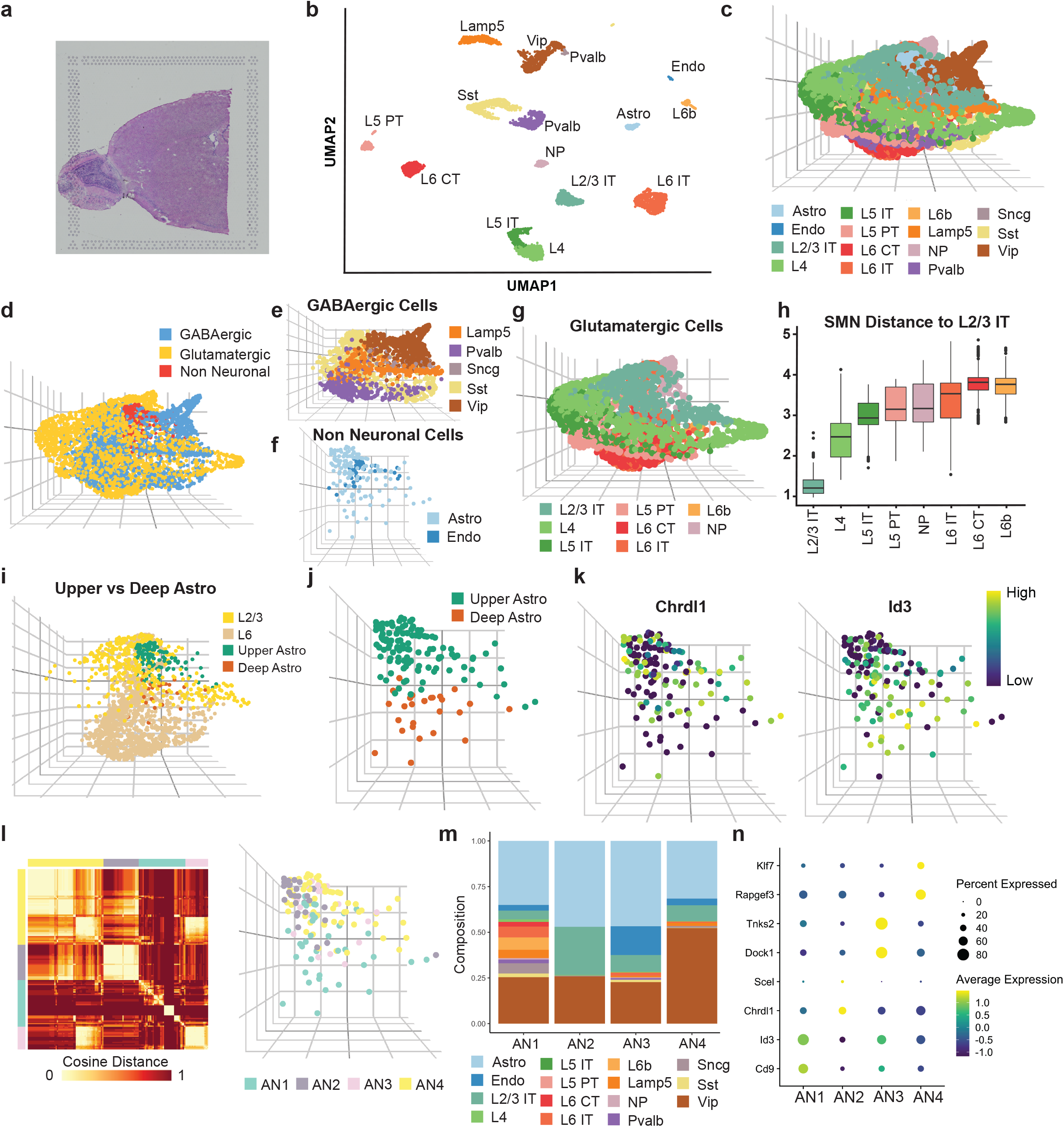
scHolography reconstructs the spatial organization of mouse brain. **a,** H&E image of an anterior brain sample for ST. **b,** UMAP plot of scRNAseq data from mouse brain. **c,** 3D visualization of spatial reconstruction of mouse brain by scHolography. **d,** GABAergic neurons, glutamatergic neurons and non-neuronal cells are visualized in the reconstructed mouse brain in 3D. **e,** Subtypes of GABAergic neurons are visualized in the reconstructed mouse brain in 3D. **f,** Non-neuronal cells, including astrocytes and endothelial cells, are visualized in the reconstructed mouse brain in 3D. **g,** Subtypes of Glutamatergic neurons are visualized in the reconstructed mouse brain in 3D. **h,** SMN distances of distinct glutamatergic neurons to L2/3 IT cells (L2/3 IT n = 353; L4 n = 489; L5 IT n = 270; L5 PT n = 188; NP n = 132; L6 IT n = 671; L6 CT n = 344; L6b n = 100). Boxplots show the median with interquartile ranges (IQRs) and whiskers extend to 1.5× IQR from the box. **i,** L2/3 (L2/3 IT) and L6 (L6 IT, L6 CT, L6b) Glutamatergic neurons are visualized together with upper and deep astrocytes. **j,** scHolography 3D plot of upper and deep astrocytes. **k,** scHolography 3D feature plot of upper (*Chrdl1*) and deep (*Id3*) astrocyte with corresponding marker genes. **l,** Spatial cell neighborhood analysis for astrocytes. Four distinct neighborhoods AN1-4 are identified based on the similarity of the first-degree neighbor cell composition (left panel). Four astrocyte spatial neighborhoods are visualized with a scHolography 3D plot (right panel). **m,** First-degree neighbor composition plot for four astrocyte spatial neighborhoods. **n,** Feature dot plot of enriched genes in each astrocyte spatial neighborhood (Mann Whitney Wilcoxon test, p<0.05).

To illustrate spatial heterogeneity within a transcriptionally defined cell type, we focused our analysis on astrocytes. A recent study, based on smFISH, has identified markers for different astrocyte layers in different cortical regions^25^. For example, *Chrdl1* expression was peaked in upper astrocytes close to L2-4 layers and *Id3* expression was peaked in deep astrocytes close to L6 layer^25^. Indeed, scHolography reconstructed mouse brain recapitulated not only the layered localization of L2/3 and L6 glutamatergic neurons but also the locations of upper and deep astrocytes (Fig. 5i and Extended Data fig. 6b). Notably, the spatial gradients of *Chrdl1 and Id3* expression in astrocytes were also recapitulated in the reconstructed brain (Fig. 5j-k and Extended Data fig. 6c). In addition to these individual gene markers, we calculated gene expression scores for the upper and deep astrocytes as a global marker for spatial astrocyte heterogeneity, and this result validated our classification of upper and deep astrocytes (Extended Data fig. 6d).

To further interrogate whether microenvironment plays a role in astrocyte heterogeneity, we performed a spatial neighborhood analysis for astrocytes. We clustered astrocytes based on their first-degree neighbor cell composition and identified 4 distinct spatial neighborhoods (AN1-4) (Fig. 5l). Closer inspection of the spatial neighbor profiles revealed that AN1 neighbors were enriched for L6 IT, L6 CT, L6b, Sncg, and Lamp5 cells, and thus these AN1 cells were related to deep astrocytes. AN2 neighbors were enriched for astrocytes and L2/3 IT cells, and thus these AN2 cells were related to upper astrocytes. AN3 neighbors were enriched for endothelial cells, and AN4 neighbors were enriched for Vip cells (Fig. 5m and Extended Data table 4). Differential gene expression analysis of each spatial astrocyte cluster further corroborated the spatial heterogeneity of astrocytes (Fig. 5n). Consistent with experimental findings, AN1 astrocytes were enriched for the deep marker, *Id3*; AN2 astrocytes were enriched for the upper marker, *Chrdl1*. Interestingly, AN3 astrocytes showed elevated expression of *Dock1* and *Tnks2*, which are related to endothelial blood-brain barrier maintenance function through WNT signaling^26^, consistent with their proximity to endothelial cells. AN4 astrocytes were differentially expressed *Rapgef3* and *Klf7*, which are associated with the Vip regulation of astrocytes^27^. Collectively, these spatial-based analyses highlight the utility of scHolography for not only faithful reconstruction of a highly complex tissue in 3D but also the identification of spatially relevant cell type clustering and gene expression pattern analysis.

## Discussion

In this study, we have provided a new computational solution to spatial transcriptomics, which defines the spatial identity of single cells, generates a neural network-based T2S projection for 3D tissue reconstruction and determines spatial cell heterogeneity. The limitation of using 2D coordinates to describe spatial identity is that the location of each pixel is determined independently by an “observer”. Thus, the interconnectedness of cell organization patterns within a tissue is not captured. As a result, the use of 2D coordinates does not accurately reflect the complex spatial organization of cells within a tissue. In contrast to using 2D coordinates, scHolography uses an inter-pixel distance matrix to describe the spatial identity of cells within a tissue. This approach relies on all pixels in the tissue to define the spatial identity of any given pixel, which preserves important information about the organization of the tissue. Additionally, the high-dimensional inter-pixel distance matrix used by scHolography enables the use of neural networks and deep learning to create an accurate projection of a cell’s transcriptome onto its spatial location. Interestingly, the T2S projection learned from low-resolution ST data is applicable to scRNAseq data without any cell-type deconvolution and, in combination with stable matching neighbor assignment, successfully reconstructs 3D tissue organization of relatively simple tissues such as human foreskin and complex tissues such as human skin cancer and mouse brain. These results show that there is a connection between a cell’s transcriptome and tissue organization, which can be revealed through the use of scHolography. Improvement in the spatial resolution and joint learning from multiple ST datasets from the same tissue should further enhance the accuracy of deep learning and reconstruction. Overall, scHolography permits the study of the effects of genetic and epigenetic perturbations on the spatial organization of cells within a tissue. The genetic information encoded in the genome determines not only a cell’s state but also the architecture of tissues and organisms. Through the use of scHolography, this can provide insights into how changes in gene expression can alter the structure of tissues or organisms. These studies have the potential to uncover new paradigms in cell-cell communication and tissue organization during development, wound healing, aging and disease.

## Methods

### The scHolography workflow

#### Step 1: Data Preparation

scHolography takes ST and SC expression data and ST 2D spatial registration data as input. scHolography first integrates the ST and SC expression data with the Seurat reference-based integration method^28^. From integrated data, scHolography obtains matrices *X_p,q_* and *Y_c,q_* where *X* are the top expression principal components (default= 32) for SC data and *Y* are the top expression principal components (default= 32) for ST data. Next, for 2D spatial registration data associated with ST data, scHolography calculates pairwise Euclidean distance matrix *D_p,p_* between spatial spots. Top *d* principal components (default =32) are then found for the distance matrix *D*, and we rename the principal components as spatial-information components (SICs). The SIC matrix is denoted as 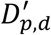.

#### Step 2: Neural Network Training

scHolography trains a neural network with *X_p,q_* as the predictor matrix and 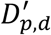 as the predicting target. The neural network functions are powered by the Keras package^29^ and have the following architecture:

| Name | Operation | Number of Features | Dropout | Batch Normalization | Activation | Input |
| --- | --- | --- | --- | --- | --- | --- |
| input | - | 32 | X | X | - | - |
| FC-1 | FC | 32 | 0.2 | √ | Leaky ReLU | input |
| FC-2 | FC | 32 | 0.2 | √ | Leaky ReLU | FC-1 |
| FC-3 | FC | 8 | 0.2 | √ | Leaky ReLU | FC-2 |
| FC-4 | FC | 32 | 0.2 | √ | Leaky ReLU | FC-3 |
| output | FC | 32 | X | X | ReLU | FC-4 |
| Optimizer | Adam |  | # of Epochs | 500 |  |  |
| Learning Rate | 0.001 | | $\alpha$ | 0.00005 | | |
| Leaky ReLU slope | 0.2 |  | Patience | 20 |  |  |
| Batch Size | 32 |  |  |  |  |  |

The network architecture is optimized with a bottleneck layer to compress information for fitting. The trained neural network will be applied to *Y_c,q_* to predict cell-specific spatial-information score *P_c,d_* corresponding to each previously identified SIC values. Based on the predicted score matrix *P*, scHolography calculates cell-cell distance and normalizes for individual cells to obtain an inferred cell-cell affinity matrix *A_c,c_*. Step 2 will be repeated for *n* times (default= 30) and the median of each *A_c,c_* entry will be found across repeated runs to reduce the variance of prediction. Denote the resulting affinity matrix as 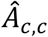 and the variance of each *A_c,c_* entry across repetitions as the learning variance matrix *M*.

#### Step 3: Spatial Neighbor Assignment

From the affinity matrix 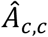, scHolography applies the Gale–Shapley algorithm to find *k* stable matching neighbors for every single cell via the MatchingR package^30^. The affinity matrix is then used as the utility for matching. Note that not all cells will be assigned *k* stable neighbors. Fewer neighbors will be assigned if there is not enough stable matching. The final stable matching results are represented in an unweighted graph. We name the graph as stable matching neighbor (SMN) graph. Once the SMN graph is determined, scHolography constructs the 3D visualization with the forced-directed Fruchterman-Reingold layout algorithm of the graph^31^. By default, the random seed is set to 60611 for all steps above.

### findDistance Function

If *a,b* are single cells within an SMN graph, we define the SMN distance between them by

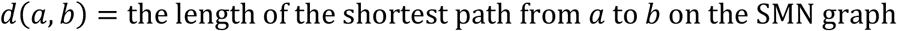

The findDistance function then enables the distance measurement of individual cells to a given cell type or cluster of cells on the SMN graph. We define the distance between a cell *x* and a cell group *A* by

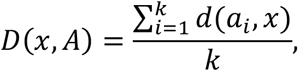

*a*_1_, …, a_k_ are the *k* nearest cells from group *A* to *x* measured by SMN distance. For default, we set *k* = 30.

### findGeneSpatialDynamics Function

The findGeneSpatialDynamics function enables the investigation of the association between spatial distribution and gene expression pattern by identifying genes with significant trends with respect to the SMN distance of cells to a reference group. Single cells in a query group Q are evaluated for their SMN distance to a reference group R. We can denote distances as *D*(*q*_1_, *R*), *D*(*q*_2_, *R*), *D*(*q*_3_, *R*) and so on. We run a Poisson regression between the expression level of each highly variable gene *i* and SMN distances to *R* of each query cell

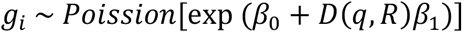

Suppose there are *n* cells in Q. *g_i_* is a vector of length *n*. *D*(*q, R*) is also a vector of length *n*

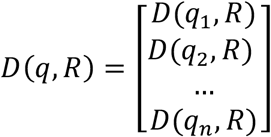

Genes are then ordered by z values from Poisson regression. Genes with negative values are considered to have proximal trends in space toward the reference group, while genes with positive z values are considered for distal trends.

### findSpatialNeighborhood Function

The findSpatialNeighborhood function aims to evaluate neighborhood cell type similarities and to define distinct spatial neighborhoods. The first-degree neighbors of a given cell are defined as stable-matching neighbors recalled from the scHolography inference. These neighbors have a direct edge to the given cell on the SMN graph. The first-degree neighborhoods are then evaluated by their composition of cell types or other given annotations. Here we define a metric named cell type frequency-inverse cell frequency (CTF-ICF). CTF-ICF inherits the idea of term frequency–inverse document frequency method for document clustering in text mining. Assume there are in *m* cell types, and there are *n* cells selected for to find neighborhoods. We first create an *m* by *n* cell type frequency (CTF) matrix *C* to count how many cells from each of the *m* cell types are present in the first-degree neighbors of *n* single cells. With the textmineR package^5^, ICF for the cell type *i* is then calculated as

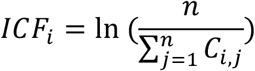

We use ICF to weigh the original CTF matrix to get the final CTF-ICF matrix 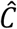

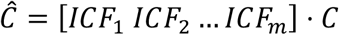

We calculated pairwise cosine similarity between cells from 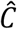, and calculated cosine distance as

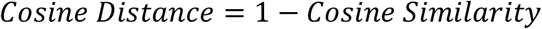

Finally, we define distinct neighborhoods by conducting hierarchical clustering on the cosine distance matrix. The number of distinct neighborhoods for clustering is optimized with the silhouette coefficient. Significant neighbor-cell types for each spatial neighborhood are identified using the one-sided Wilcoxon test with p-values<0.05.

### scHolographyNeighborCompPlot Function

The scHolographyNeighborCompPlot function plots the first-degree neighbor composition with respect to a given annotation. The function also identifies enriched neighbor types for query cells with significance levels using the Wilcoxon test.

### Human foreskin sample collection and sequencing

Neonatal foreskins from Donors 1 and 2 were collected as discarded, deidentified tissue under IRB protocol #STU00009443 of the Northwestern University Skin Biology and Diseases Resource-based Center. Donor 1 sample was punched by an 8mm punch and embedded in the sagittal direction into an FFPE block by SBDRC.

For the scRNA-seq experiment, fresh human foreskin specimens from Donor 2 were cut into 4 mm x 4 mm pieces. The dermal fat layer was trimmed off from the bottom. Then the skin was floated on 2 mL of dispase in a 6-well plate and incubated at 37 °C for 1 hour. The epidermis was separated from the dermis and trypsinized for 12 minutes at 37 °C to get the epidermal single-cell suspension. For the dermis part, it was further cut into smaller pieces, then incubated with 0.25% collagenase I in 2 mL HBSS for 1 hour at 37 °C. Collagenase-treated pieces were trypsinized for 10 minutes at 37 °C. The tissue was then dissociated by pipetting and single-cell suspension was obtained. Epidermal and dermal cells were combined at a 1:1 ratio and used as scRNA-seq input materials. The Single-Cell Chromium 3’ v3 kit from 10x Genomics was used for single-cell library preparation. Final scRNA-seq libraries were sequenced on an Illumina NovaSeq-6000 system.

The Cell Ranger v.6.0.0 was applied to align reads to the human reference GRCh38 (GENCODE v32/Ensembl 98), and a gene expression matrix was obtained. The Seurat package v4 was used for data processing and visualization, and the default settings were applied unless otherwise noted. Cells with fewer than 200 or more than 7000 unique feature counts were filtered. Besides, cells with more than 15% of mitochondrial counts were also filtered. The normalization was performed by sctransform^32^. Variable genes were found with the FindVariableFeatures function and PCA was conducted by RunPCA. The top 30 PCs were selected with ElbowPlot for downstream analyses. Cell clusters were identified by FindNeighbors and FindCluster functions at a resolution of 0.5. RunUMAP was used for 2D visualization. DE genes were identified by the FindAllMarkers function and the top DE genes for each cluster were considered for cell identity annotation.

For ST experiments, RNA quality was first checked for the sample. Total RNA was isolated from a 20um Donor 1 FFPE block section using Qiagen RNeasy FFPE Kit following the manufacturer’s instructions. RNA quality was evaluated using the DV200 assay on Agilent Bioanalyzer. The sample was used for library preparation after confirming the quality of RNA is desired based on DV200 (DV200 > 50%; DV200 = proportion of RNA fragments with >200 nucleotides in length).

Two 5um sections in serial were sliced from Donor 1 FFPE block, placed on 10X Genomics Visium Spatial Gene Expression Slide v1, deparaffinized, and H&E stained under the manufacturer’s protocol. Two samples were placed on A1 and B1 capturing regions, respectively. Brightfield images were acquired at 20x magnification using a Nikon Ti2 widefield microscope system for 2 hours. Images were processed with the Nikon NIS-elements software. The samples were then decrosslinked, and the human whole transcriptome probe panel was hybridized to the RNA from the decrosslinked tissue. Next, probes were ligated, released from the tissue, extended, and indexed. All these steps followed the manufacturer’s instructions. For library construction, 17 cycles of sample index PCR were performed.

Final ST libraries were sequenced on an Illumina NovaSeq-6000 system. The Space Ranger v.1.3.1 was applied to align reads to the human reference GRCh38 (GENCODE v32/Ensembl 98). The Seurat package v4 was again used for data processing and visualization, and the default settings were applied unless otherwise noted. The normalization was performed by sctransform^32^. Variable genes were found with the FindVariableFeatures function and PCA was conducted by RunPCA. The top 32 PCs were selected for downstream analyses. Pixel clusters were identified by FindNeighbors and FindCluster functions at a resolution of 0.5. RunUMAP was used for 2D visualization. DE genes were identified by the FindAllMarkers function.

### Human foreskin data analysis

Donor 2 scRNA-seq data were reconstructed by scHolography using Donor 1 slice 1 ST data as the reference. Default scHolography settings were used. For benchmarking, we use Donor 1 slice 2 ST data as a testing dataset. We reconstructed Donor 1 slice 2 expression data using Donor 1 slice 1 ST as the reference. We compared the reconstruction results of Donor 1 slice 2 to its true spatial registration information. We computed the pixel-wise SMN distance matrix *D_scHolography_* and pixel-wise Visium spatial registration Euclidean distance *D_Visium_* of Donor 1 slice 2. Two metrics, Spearman correlation and Jaccard similarity, were calculated for *D_scHolography_* and *D_Visium_* to evaluate global and local prediction accuracy, respectively. Specifically, the two metrics for pixel *i* were defined as

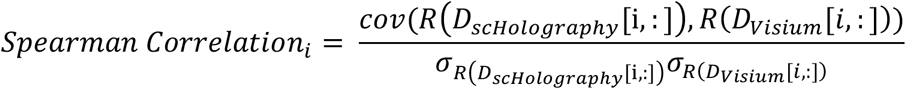

Where *R* is ranks and *σ_R_* is the standard deviation of the ranks.

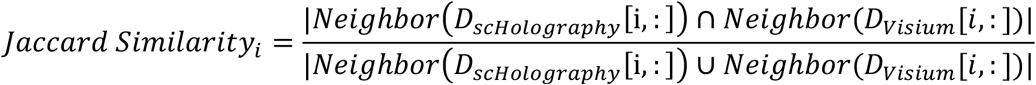

Where *Neighbor* is a set for 30 nearest neighbors for pixel *i* under either *D_scHolography_* or *D_Visium_*.

For comparison, CellTrek and Seurat-SrtCT predictions were also performed on Donor 2 scRNA-seq data and Donor 1 slice 2 ST data using Donor 1 slice 1 ST data as the reference. We rank Cell Trek with following parameters: intp_pnt=999, nPCs=30, ntree=1000, dist_thresh=999, top_spot=1, spot_n=999, repel_r=20 with 20 iterations. This setting aimed to reduce the number of unmapped cells for a fair comparison. Default settings of Seurat label transfer were used with dims=1:30. CellTrek and Seurat-SrtCT inferred Donor 2 scRNA-seq data were visualized in 3D by first computing cell-wise inferred spatial Euclidean distance matrices *D_CellTrek_* and *D_Seurat_*. The distance matrices were then employed as utilities to be fed into scHolography for SMN graph construction and 3D visualization.

The CellChat analysis^13^ was performed to dissect ligand-receptor interactions for suprabasal and basal cells in Donor 2 scRNA-seq data with default settings. Unless otherwise noticed, all differential gene expression analyses for this paper used the Wilcoxon test that is powered by FindAllMarkers and FindMarkers functions of Seurat.

### Human cSCC data acquisition and analysis

The filtered gene count matrices of the human cSCC 3’ scRNA-seq data were downloaded from GEO (GSE144240), and the cell types were annotated based on the level 2 cell types from the original study^7^. Data were subsetted to keep only Patient 6 data. The keratinocyte cluster without specific keratinocyte state annotations and the multiplet cluster were excluded from downstream processing. The human cSCC ST data was also downloaded from GEO (GSE144240). Only two replicates from Patient 6 were processed. The analysis and visualization were handled by automated processing and integration steps of scHolography workflows built upon Seurat (SCTransform normalization, nPCtoUse=32, FindCluster.resolution=0.5).

scHolography prediction of cSCC scRNA-seq data was performed using Patient 6 replicate 1 ST data as the reference. For validation and benchmarking purpose, Patient 6 replicate 2 ST expression data was used. The scHolography, CellTrek, and Seurat-SrtCT predictions of Patient 6 replicate 2 were then compared with the true Patient 6 replicate 2 spatial registration results from ST. The parameters for the three methods were the same as the previous human skin analysis. Spearman correlation and Jaccard similarity were also calculated as previously described.

### Mouse brain data acquisition and analysis

The mouse brain scRNA-seq and ST data were downloaded from the CellTrek website^6^ (https://github.com/navinlabcode/CellTrek). Only the frontal cortex region of ST data was processed. The cell types were annotated based on the cell type from the original study^8^. The analysis and visualization were handled by automated processing and integration steps of scHolography workflows built upon Seurat (SCTransform normalization, nPCtoUse=32, FindCluster.resolution=0.5).

scHolography prediction of mouse brain scRNA-seq data was performed using the mouse brain ST data as the reference. For the astrocyte analysis, we calculated the SMN distances of all astrocyte cells to L2/3 (L2/3 IT) and L6 (L6 IT, L6 CT, and L6b) using the findDistance function. Astrocytes were classified as Upper Astro if they were closer to L2/3 in terms of SMN distance. Otherwise, astrocytes were classified as Deep Astro. Layer astrocyte markers were obtained from a previous study^25^. Upper Layer and Deep Layer scoring were performed using the Seurat AddModuleScore function with parameters used in the developer’s tutorial.

## Data availability

The human foreskin scRNA-seq and ST data were submitted to the Gene Expression Omnibus (GEO): GSE220573.

## Code availability

scHolography code and documentation are available at: https://github.com/YiLab-SC/scHolography.

## Acknowledgments

We thank members of the Yi and Braun laboratories for suggestion and discussion. We thank the NU-SBDRC Skin Tissue Engineering and Morphology Core, supported by National Institute of Health Grant P30AR075049, for assistance for human foreskin sample collection and preparation. This work was supported by National Institute of Health Grant R01AR066703, R01AR071435, R01AR081103 and R01HD107841 (RY) and a pilot grant from Northwestern University Skin Biology and Diseases Resource-based Center grant P30AR075049. R.B. was supported by NSF 1764421-01 and SFAR 597491-RWC01.

## Contributions

R.Y. and Y.F. conceived the study. Y.F. performed human foreskin ST experiments with assistance from A.D., Y.F. performed method development and computational analysis. D.W. generated human foreskin scRNAseq dataset. R.B. supervised computational method development and analysis. R.Y. supervised the study and wrote the manuscript together with Y.F. with input from all authors.

---

**Extended Data Figure 1.**
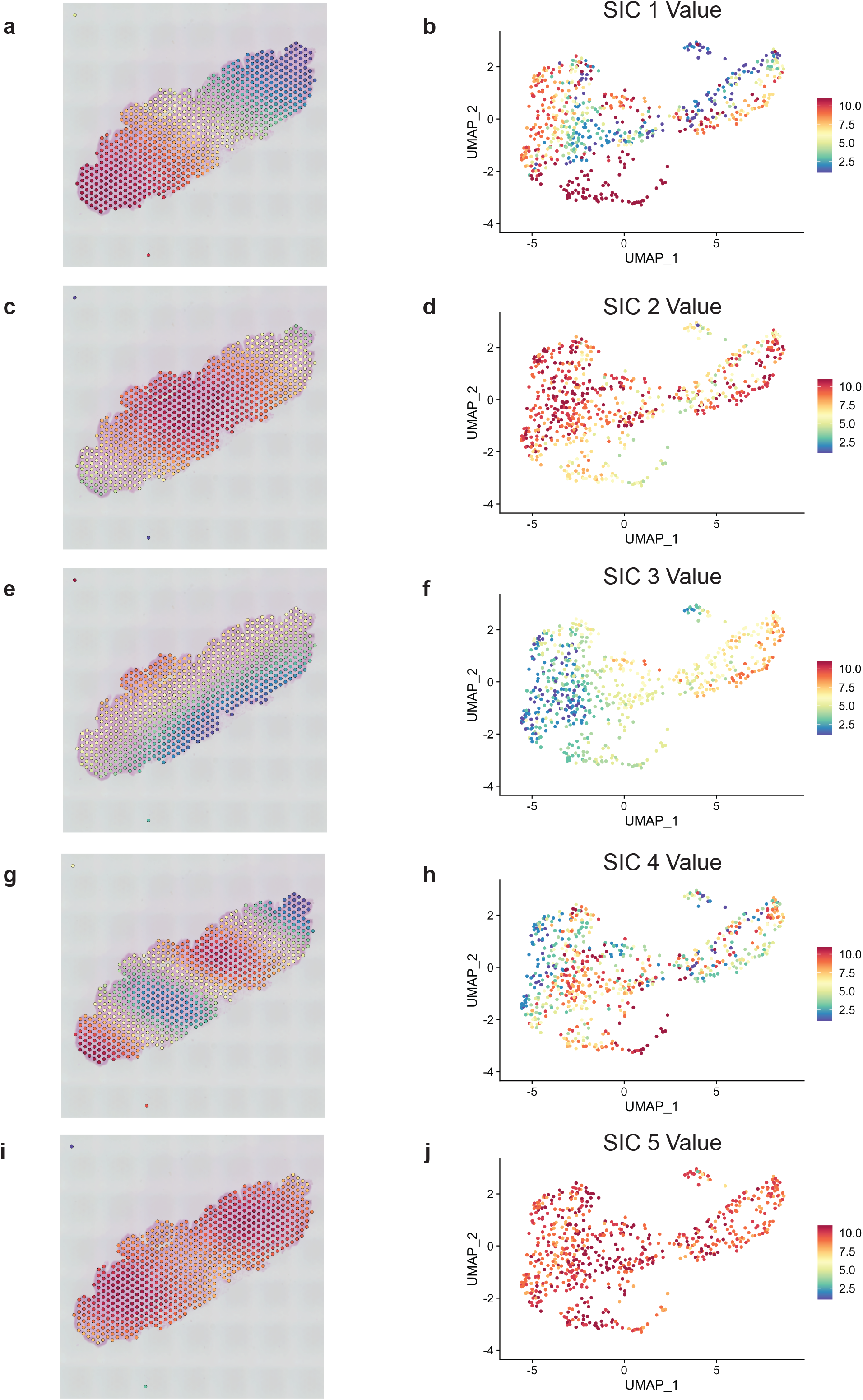
Spatial relevance of SICs. Top five SIC values of ST data from Donor 1 slice 1 on spatial image (left panels) or expression UMAPs (right panels) are plotted to demonstrate the pattern correlation between spatial organization and expression profiles. **a-b,** SIC 1 values. **c-d,** SIC 2 values. **e-f,** SIC3 values. **g-h,** SIC 4 values. **i-j,** SIC 5 values.

**Extended Data Figure 2.**
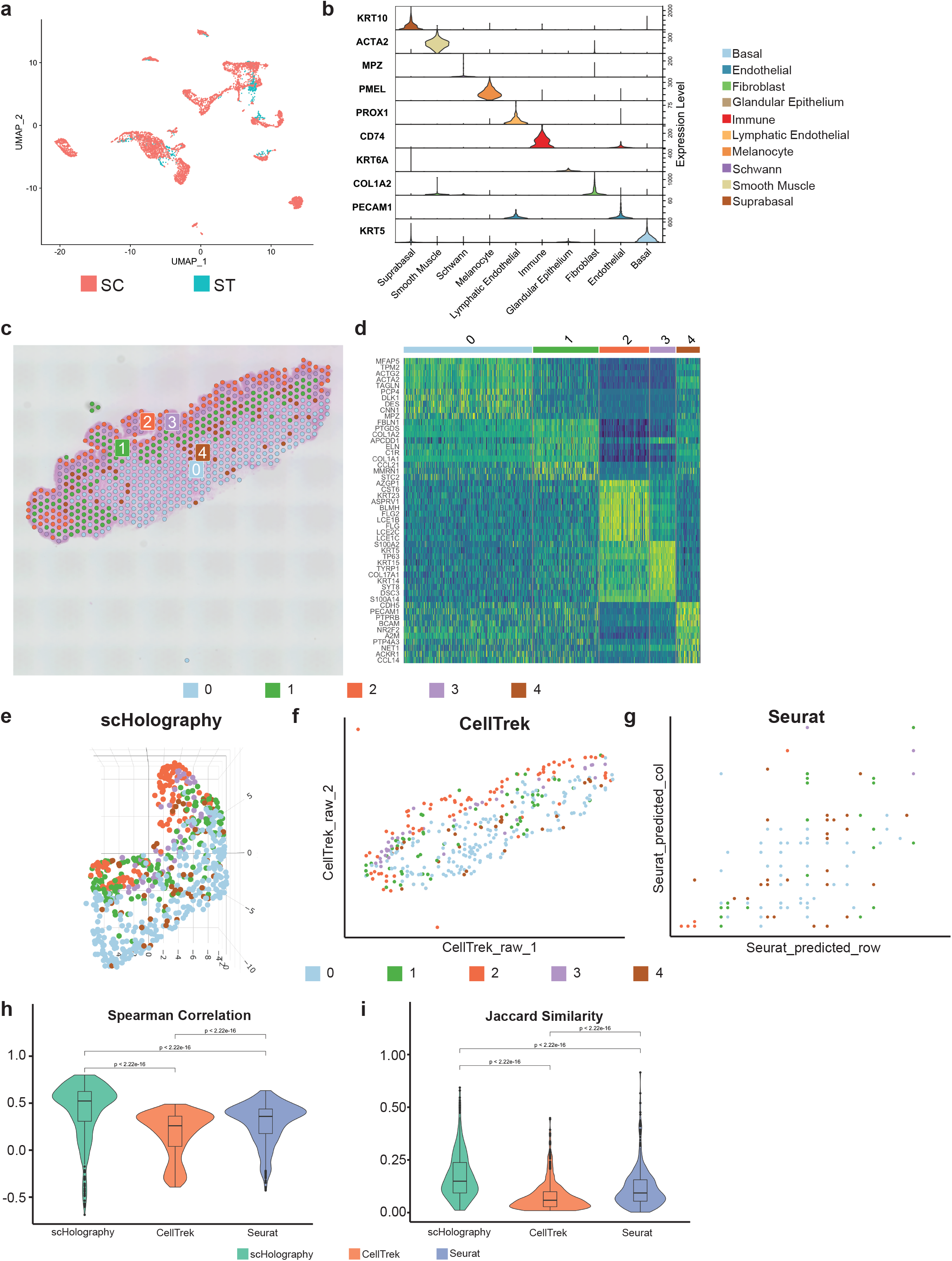
Benchmarking of scHolography with current methods. **a,** UMAP plot of scRNAseq and ST data integration. **b,** Identification of major cell types in human foreskin samples by cell lineage markers. *KRT10*, suprabasal epithelial cells; *ACTA2*, smooth muscle cells.; *MPZ*, Schwann cells; *PMEL*, melanocytes; *PROX1*, lymphatic endothelial cells; *CD74*, immune cells; *KRT6A*, glandular epithelial cells; *COL1A2*, fibroblasts; *PECAM1*, endothelial cells; *KRT5*, basal epithelial cells. **c,** Spatial spot plot of ST data from Donor 1 slice 2. Five clusters are identified. **d,** Expression heatmap of top ten markers of each of the five clusters from Donor 1 slice 2 ST data (Mann Whitney Wilcoxon test, p<0.05). **e,** 3D plot of the reconstruction result from Donor 1 slice 2 ST data by scHolography. **f,** Dot plot of the reconstruction result from Donor 1 slice 2 ST data by Celltrek. **g,** Dot plot of the reconstruction result from Donor 1 slice 2 ST data by Seurat-SrtCT. **h,** Violin plot of Spearman correlation for the comparison of prediction accuracy by scHolography, CellTrek, and Seurat-SrtCT. One-sided Wilcoxon tests are performed to determine statistical significance. **i,** Violin plot of Jaccard similarity for the comparison of prediction accuracy by scHolography, CellTrek, and Seurat-SrtCT.

**Extended Data Figure 3.**
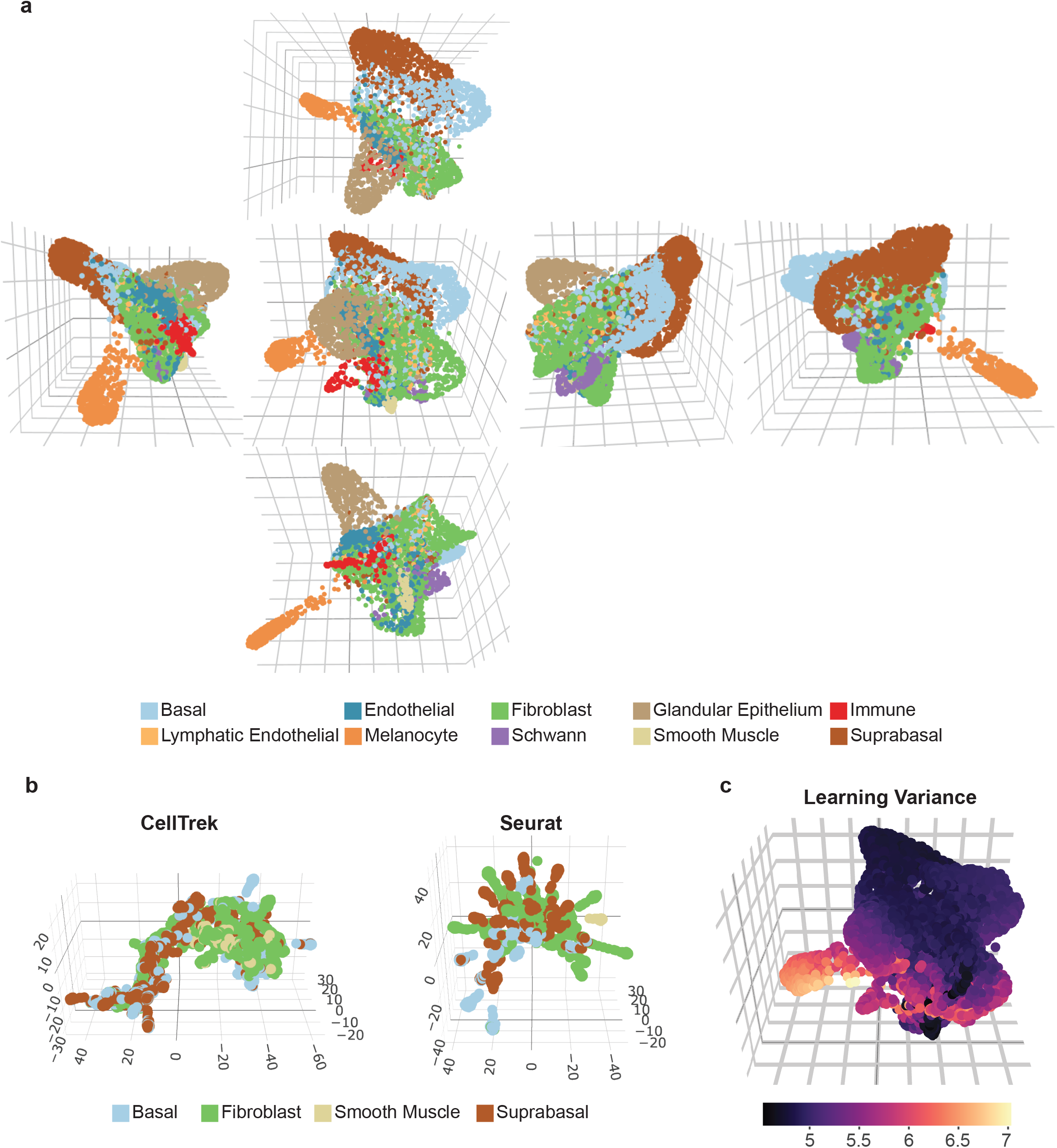
Three-dimensional reconstruction of human foreskin. **a,** Cubemap of single-cell human foreskin reconstruction by scHolography. The cubemap includes perspective snapshots from the top (row 1), left (row 2, column 1), front (row 2, column 2), right (row 2, column 3), back (row 2, column 4), and bottom (row 3) of scHolography reconstruction. **b,** 3D plot of CellTrek (left) mapping and Seurat-SrtCT (right) mapping results. The spatial cell-cell distance matrices are calculated from CellTrek and Seurat-SrtCT mapping results and fed into scHolography for SMN graph construction and 3D visualization. **c,** 3D feature plot of learning variance of each cell by scHolography.

**Extended Data Figure 4.**
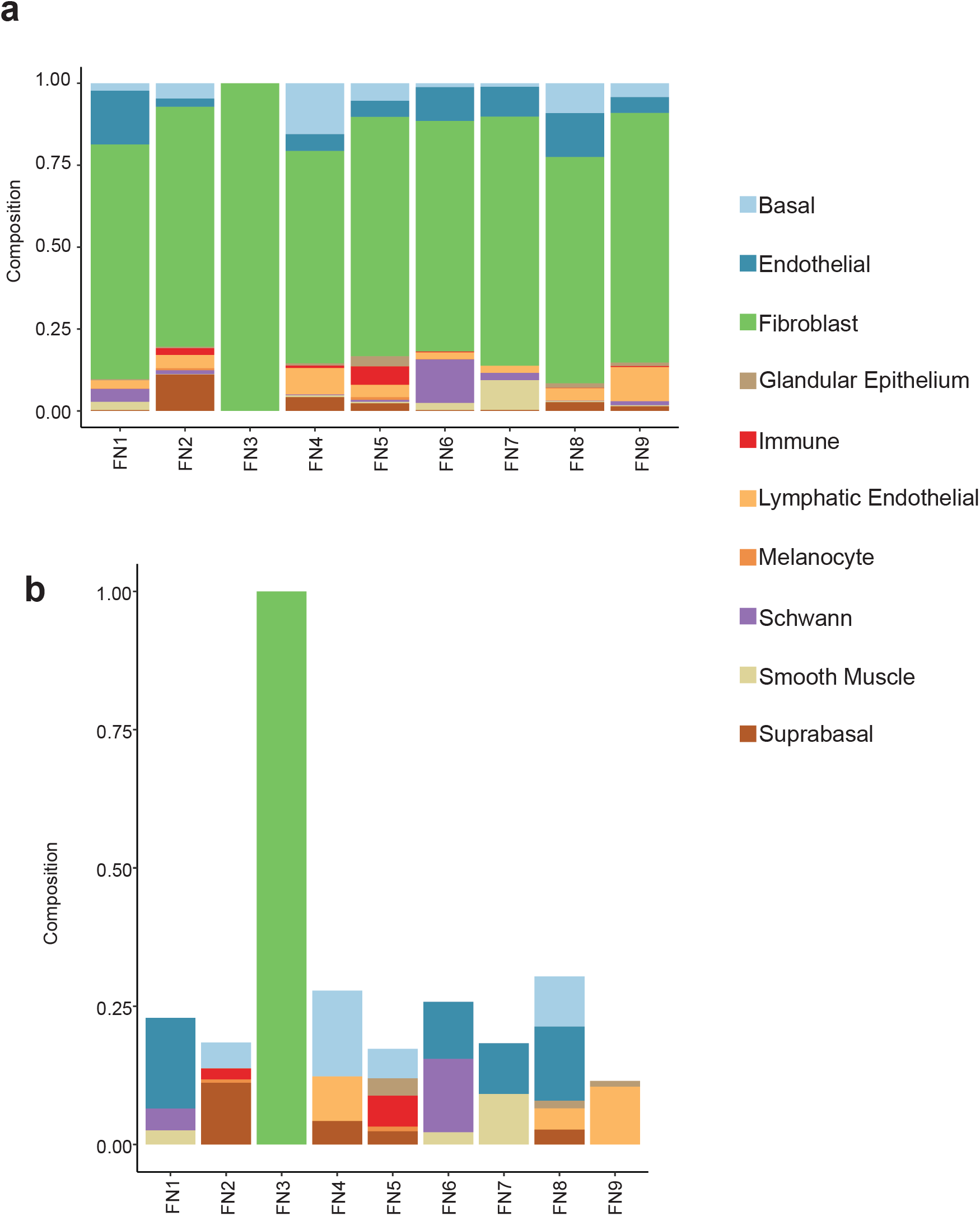
Spatial neighborhoods of fibroblasts in human foreskin. **a.** First-degree neighbor composition plot of FN1-9. **b.** First-degree neighbor composition plot for significantly enriched neighboring cell types of FN1-9 (One-sided Mann Whitney Wilcoxon test, p<0.05).

**Extended Data Figure 5.**
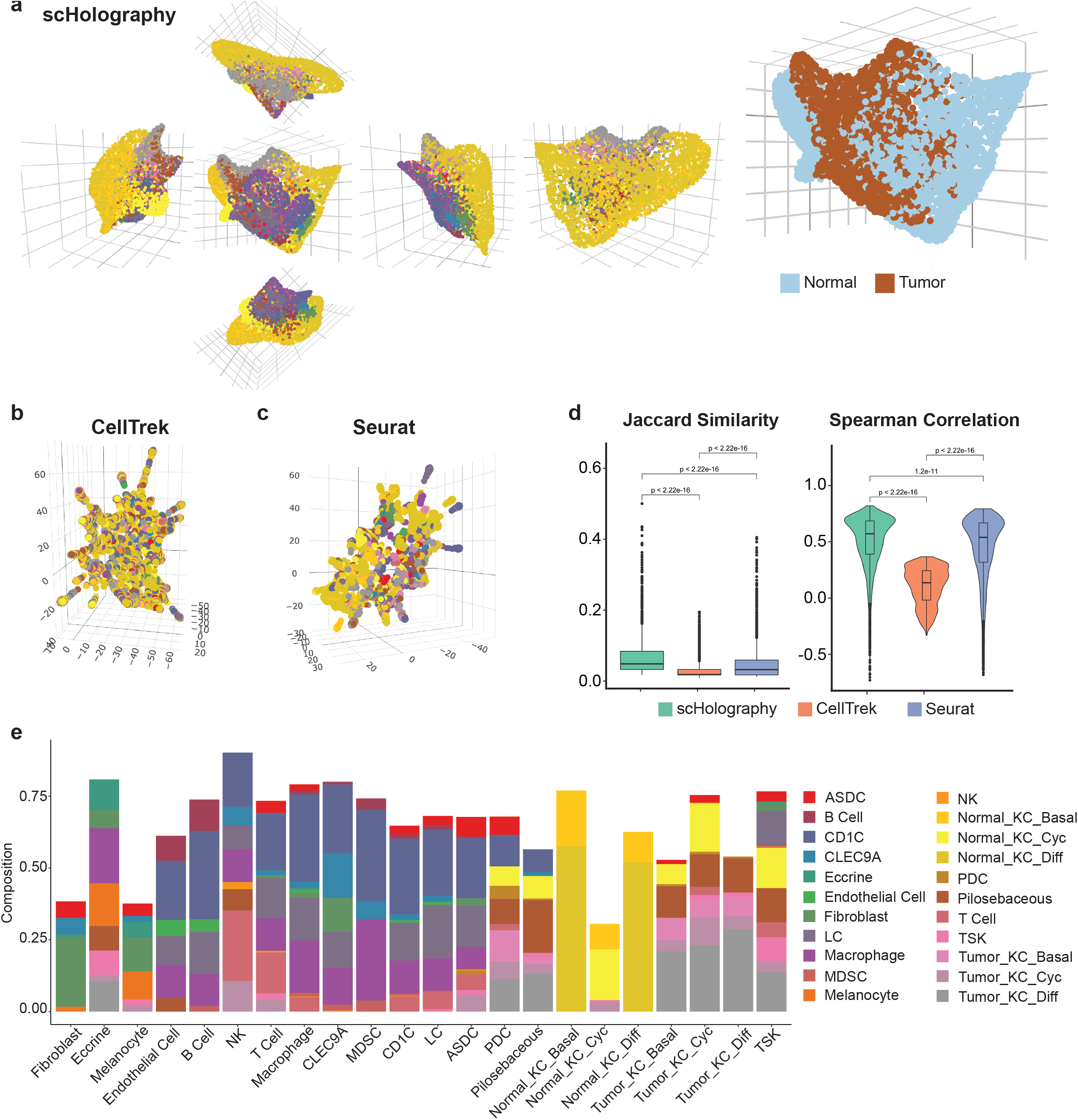
Benchmarking of scHolography and spatial neighborhood profiles of human cSCC. **a,** Cubemap of single-cell cSCC reconstruction by scHolography (left) and scHolography 3D plot of cSCC spatial reconstruction colored by normal and diseased conditions (right). **b-c,** 3D plot of CellTrek (**b**) mapping and Seurat-SrtCT (**c**) mapping results. **d,** Violin plots of Jaccard similarity (left) and Spearman correlation (right) for the comparison of prediction accuracy of scHolography, CellTrek, and Seurat-SrtCT. One-sided Wilcoxon tests are performed to determine statistical significance. **e,** First-degree neighbor composition plot for significantly enriched neighboring cell types within cSCC (One-sided Mann Whitney Wilcoxon test, p<0.05).

**Extended Data Figure 6.**
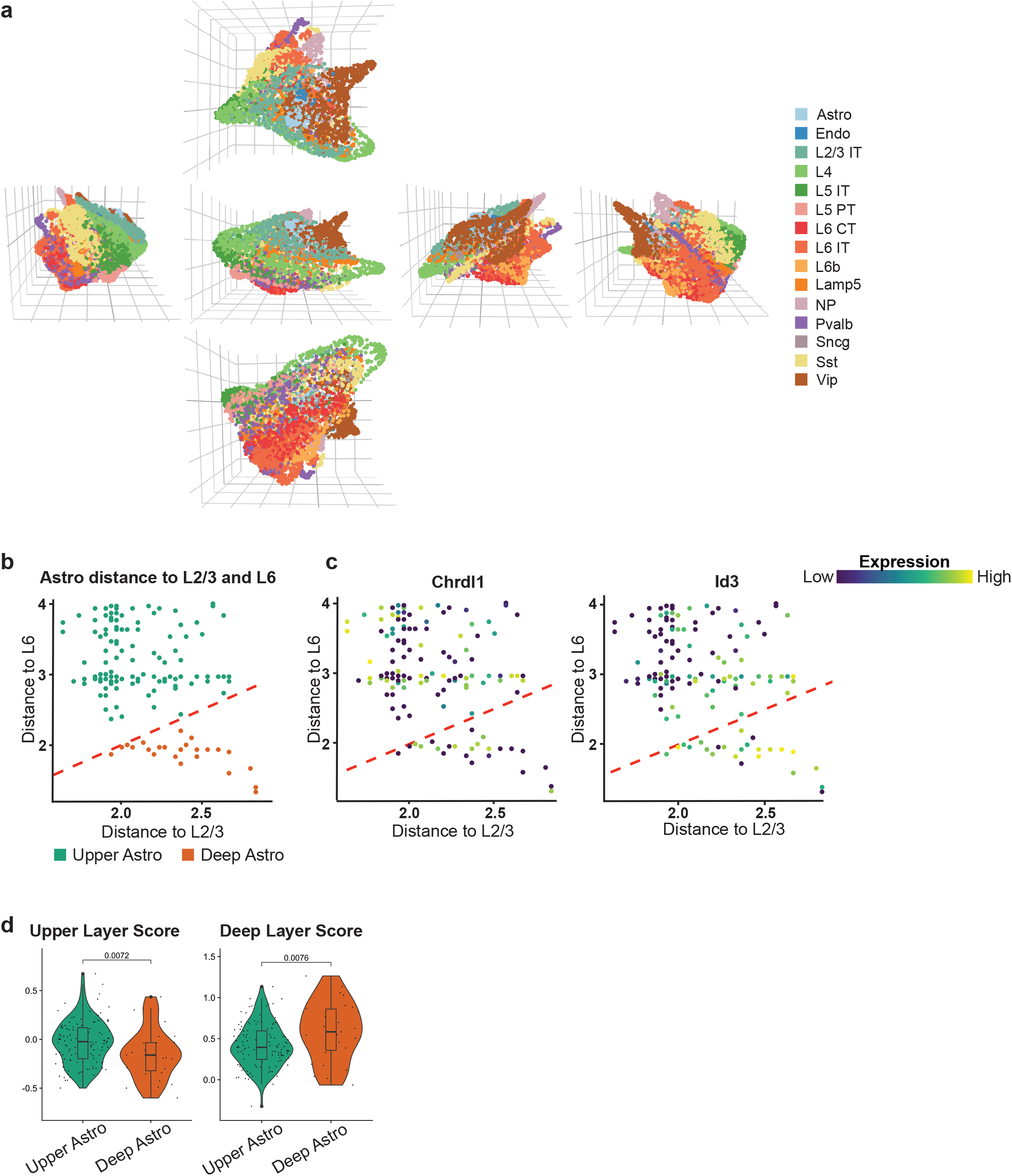
Three-dimensional reconstruction of mouse brain and spatial neighborhood analysis for astrocytes. **a,** Cubemap of single-cell mouse brain reconstruction by scHolography. **b,** Distance plot of astrocytes to L2/3 and L6 colored by upper and deep astrocyte layers. **c,** Distance plot of astrocytes to L2/3 and L6 colored by *Chrdl1* (left, upper astrocyte marker) and *Id3* (right, deep astrocyte marker) expression. **d,** Violin plots of upper layer score (left) and deep layer score (right) in upper and deep astrocyte layers. One-sided Wilcoxon tests are performed to determine statistical significance.

Extended Data table 1. First-degree neighbor composition by cell type in human foreskin.

Extended Data table 2. Spatial neighborhood composition for fibroblasts in human foreskin.

Extended Data table 3. First-degree neighbor composition by cell type in human cSCC.

Extended Data table 4. Spatial neighborhood composition for astrocytes in mouse brain.

